# Evidence accumulation under uncertainty – a neural marker of emerging choice and urgency

**DOI:** 10.1101/2020.06.30.179622

**Authors:** Elisabeth Parés-Pujolràs, Eoin Travers, Yoana Ahmetoglu, Patrick Haggard

## Abstract

To interact meaningfully with its environment, an agent must integrate external information with its own internal states. However, information about the environment is often noisy. In our task participants had to monitor a stream of discrete visual stimuli over time and decide whether or not to act, on the basis of either strong or weak evidence. We found that the classic P3 event-related potential evoked by sequential evidence items item of evidence tracked decision-making processes and encoded participants’ choice, both when evidence was strong and when it was weak. We also found that the readiness potential, a classic marker of self-paced actions, was observed preceding all actions - even when those were strongly driven by external evidence. Computational modelling showed that both neural dynamics and behavioural results can be explained by a combination of (a) competition between mutually inhibiting accumulators for the two categorical choice outcomes, and (b) a context-dependent urgency signal.

## 1. Introduction

To interact meaningfully with its environment, an agent must integrate external information with its own internal states. Human agents often face decisions where there are no compelling external or internal reasons to decide one way rather than another. Since there is only a limited amount of time available to make any given decision, successful behaviour requires the capacity to break the symmetry between alternatives and pick one of them. How do we decide what to do when the available information is ambiguous?

A traditional approach to study this question in voluntary action has been to study the endogenous brain mechanisms that enable human agents to spontaneously trigger action in the absence of any externally relevant input (e.g. Libet et al., 1983). Yet, most voluntary actions are typically *informed* by the environment, even when they are not determined by it. In many daily-life scenarios agents seek external information to help them decide, but often such information remains inconclusive. In those cases, evidence-driven decisions can effectively become ‘perceptual guesses’ (Bode et al., 2012). Interestingly, disambiguation of perceptual stimuli shares similarities with spontaneous action initiation: both types of situations require the ability to make a decision and initiate action in the absence of compelling exogenous triggers. Which brain mechanisms enable decision and action in such contexts?

Perceptual decision-making is often described as a sequential sampling process, where evidence is continuously sampled and accumulated, triggering a decision or action once a given threshold is reached (Forstmann et al., 2016). It has been suggested that symmetry-breaking mechanisms enabling action in the absence of clear exogenous information may rely on noisy accumulation processes (Bode et al., 2013; Schurger et al., 2012). If no strong evidence is present, small fluctuations in stimulus properties (Charles & Haggard, 2020) or intrinsic neural activity (Deco & Romo, 2008; Glimcher, 2005; Maye et al., 2007) can introduce asymmetries that eventually determine decisions. Further, it has been suggested that a time-varying urgency signal may facilitate the termination of decision-making processes and potentially action execution by adjusting the accuracy criterion in a context-dependent manner (Churchland et al., 2008; Ditterich, 2006; Thura, 2020). Urgency has often been conceptualised as a *multiplicative* gain modulation of the decision-relevant input (Cisek et al., 2009; Ditterich, 2006; Murphy et al., 2016; Thura & Cisek, 2014), but also as an *additive* signal that increases the firing rate across neuronal populations (Churchland et al., 2008). In this study, we aimed to investigate the neural correlates of these potential symmetry-breaking mechanisms at the sensory and motor levels.

At the sensory processing level, we aimed to track the evolution of category-specific decision signals to investigate when asymmetries in the processing of competing alternative options arise. Single-cell studies have shown that the firing rate of single neurons in lateral intraparietal area (LIP) build to threshold tracking decision-making processes in macaque monkeys (Kiani et al., 2008; Roitman & Shadlen, 2002; Shadlen & Newsome, 2001; Shadlen et al., 1996), and that distinct populations encode stimuli that are assigned to specific response categories in perceptual decision-making tasks (Fitzgerald et al., 2011; Freedman & Assad, 2006; Swaminathan & Freedman, 2012). Human neural correlates of such categorical processing during evidence accumulation in decision-making have not yet been described, but some clear candidate signals have been previously identified with EEG.

The P3 event-related potential has been shown to exhibit build-to-threshold dynamics, correlating with the difficulty and reaction times in perceptual response tasks (Kutas et al., 1977; Twomey et al., 2015). Further, the centro-parietal positivity (CPP), an EEG signal sharing neural sources with the P3, similarly tracks the evolution of multimodal decisions in change detection tasks (O’Connell et al., 2012; Steinemann et al., 2018) and also categorical choices about random dot motion stimuli (Kelly & O’Connell, 2013) over longer time scales. These signals may therefore provide useful neural markers of an internal decision variable. However, previous research on categorical choice using random dot motion tasks could not separately resolve the neural activity evoked by the two categories of evidence, since both leftward and rightward motion were simultaneously present. Here, we developed a new paradigm that allowed us to investigate whether P3 responses to sequential pieces of evidence could be used to track not only the general state of an internal decision variable (i.e. how close to making a decision people are), but also the relative changes in activity of the underlying category-specific neural populations (i.e. *which* decision people are going to make).

At the motor level, we were interested in the potential role of motor preparation activity as a symmetry-breaking mechanism in self-paced actions. The Readiness Potential (RP) is a slowly ramping negativity in motor areas that has been identified as a reliable neural precursor of spontaneous action (Kornhuber & Deecke 1965; Keller & Heckhausen, 1990; Khalighinejad et al., 2018; Schultze-Kraft et al., 2016; Shibasaki & Hallett, 2006; Trevena & Miller, 2002). Since this signal appears to be absent in cued-reaction tasks (Jahanshahi et al., 1995; Papa et al., 1991), it has traditionally been interpreted as a marker of endogenous action preparation and initiation. However, RP shapes can be obtained by averaging of stochastic fluctuations in the motor system in humans (Schurger et al., 2012), and spontaneous behaviour in mice (Murakami et al., 2014, 2017) is preceded by a similar build-up of activity in motor areas. Thus, it has been suggested that the RP may reflect noisy accumulation of internal noise and play a determinant role in triggering arbitrary decisions, but not deliberate ones (Maoz et al., 2019).

In this experiment, participants had to decide *whether* to make an action on the basis of extended sequences of discrete stimuli. Each discrete stimulus either favoured acting (positive evidence, +Ev), or withholding action (negative evidence, –Ev). Each trial lasted for exactly 25 s and participants were not rewarded for acting fast, so the timing of their decisions and actions was largely self-paced. Further, actions in this task were neither direct reactions to a single immediate stimulus, nor completely spontaneous and independent from the current environment. The proportion of stimuli providing positive and negative evidence varied across trials. Some *informative* trials involved easy decisions, since they showed strong net evidence in favour of acting or not acting. Other, *neutral* trials were very difficult, since the letter stream presented no net evidence in favour of either option (i.e. +Ev and –Ev stimuli were equally likely). This type of discrete stimuli are in sharp contrast with other decision-making paradigms (e.g. random-dot motion (Kelly & O’Connell, 2013), or changing contrast (O’Connell et al., 2012)), and they crucially allowed us to investigate the emergence of *choice-specific* decision signals with each successive, discrete item of evidence. Further, the comparison between action initiation signals in the externally-determined *(informative)* vs. underdetermined (*neutral*) scenarios allowed us to shed new light onto the significance of the RP.

## 2. Results

### 2.1.1. Task & design

Participants (n= 19, n = 3 excluded from EEG analysis) performed a simple perceptual decision-making task (*Figure 1a*). Each trial started with either a blue or a pink background, and a 4 Hz, 25 s letter stream containing relevant and distractor stimuli was presented. Participants were instructed to monitor the letter stream and decide whether one set of target letters (*b,d*) appeared more frequently than another set (*p,q*). Participants’ task was to make sure that the background colour matched the more frequent group of letters (i.e. if the most frequent set of targets was *bd,* the screen should be blue. If the most frequent set of targets was *pq,* the screen should be pink). If the initial colour of the screen on a given trial matched the set of targets participants perceived as most frequent, they were not required to do anything (*No-action* trials). If the initial colour of the screen on that trial did not match the most frequent set of targets, they had to press the key corresponding to the colour they wanted to switch it to (i.e. either pink or blue) with the left or right hand respectively (*Action trials*). They were allowed to switch the colour of the screen only once per trial. There was no incentive to respond fast, since trials were not terminated at the time of response and stimuli continued to appear on the screen until the end of the 25 s. At the end of each trial, participants were asked to report how confident they were that changing the colour of the screen (or not changing it) was the correct choice, using a Visual Analog Scale (VAS). If they had acted, they were also asked to estimate at what point during the trial they acted, again using a VAS.

**Figure 1.**
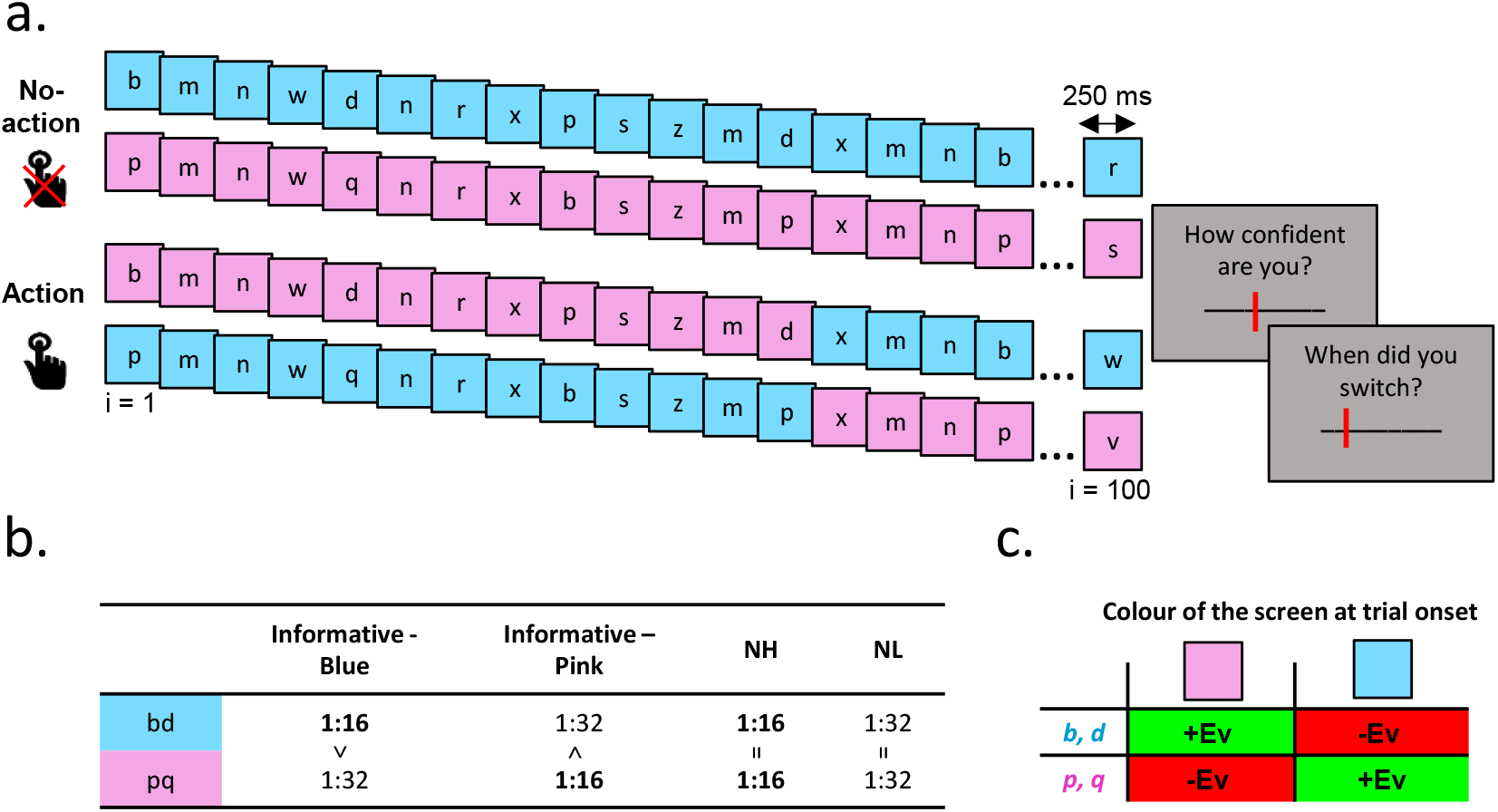
Task and experimental design. **a.** Each trial started with either a blue or a pink background. A letter stream was presented for 25 seconds (100 letters, 250 ms each), and participants had to decide which of the two target letter sets was most frequent (blue targets: ‘b’ and ‘d’, or pink targets: ‘p’ and ‘q’). Their task was to make sure that the colour of the screen matched the most frequent set of targets (e.g. if ‘p’ and ‘q’ were most frequent, the screen should be pink). Sometimes, the starting colour would already match the set of targets participants perceived as most frequent. In those cases, they were instructed to not execute any action (No-Action). In other trials, the most frequent targets did not match the given colour of the screen (Action). In those cases, they could change the colour of the screen by pressing either the ‘f’ key with their left h and to turn the screen pink, or the ‘j’ key with their right hand to turn it blue. They were only allowed to change the colour of the screen once every trial. After 25 s, the letter stream was terminated and participants were asked confidence ratings and estimates of the time at which they made a decision in Action trials only. **b.** The relative frequency of relevant targets defined both conditions. In informative conditions, the most frequent set of targets appeared twice as often as the less frequent one (once every 16 letters, or once every 32). In the neutral conditions, both sets of targets appeared equally often. In the NH condition they both appeared very often (once every 16 letters), while in the NL condition they appeared very rarely (once every 32 letters). **c.** Coding scheme. Relevant letters were coded with respect to action, given the initial colour of the screen. In trials where the screen was Blue at trial onset, pq were evidence for action (positive evidence, +Ev), and bd were evidence against action (negative evidence, –Ev).

The frequencies of the two sets of targets, together with the colour of the screen at trial onset defined four experimental conditions (*Figure 1b*). In the *Informative* conditions, the most frequent set of letters would appear twice as frequently as the other set. The stimuli were presented at a random time every 1 to 4 s (frequent stimuli) or every 1 to 8 s (infrequent stimuli). Thus, there was strong evidence either for or against action. In the *Informative anti-Action* condition, the colour of the screen at trial onset and the most frequent set of targets matched (i.e. the frequency of *pq>bd* and the screen was pink, or frequency of *pq* < *bd* and the screen was blue). Hence, participants should not press a key to change the colour of the screen. In the *Informative pro-Action* condition, the colour of the screen and the most frequent set of targets did not match (i.e. the frequency of *pq>bd* and screen was blue, or the frequency of *pq* < *bd* and the screen was pink). Therefore, participants should press a key to switch the colour of the screen.

In *Neutral* conditions, both sets of letters were presented at the same frequency. Two neutral conditions were created so that, on average, the overall number of relevant stimuli (i.e. rate) across the two neutral conditions would be equivalent to that in the informative ones. In the Neutral High (*NH*) input condition, evidence was abundant (i.e. one piece of +Ev and one piece of –Ev were presented at a random time every 1 to 4 s). In the Neutral Low (*NL*) input condition, evidence was scarce (i.e. one piece of +Ev and one piece of –Ev were presented at a random time every 1 to 8 s). In these two neutral conditions, there was no *net* evidence for or against switching in any given trial. Note, however, that participants were *not* told that in some conditions there was no net evidence. In all conditions, letters from both sets of targets were never presented within intervals shorter than 0.5 s.

Participants performed a total of 39 trials per condition. The experiment was divided in 3 blocks of 52 trials, and all conditions were randomised within blocks (13 trials per condition per block).

### 2.1.2. Behavioural results

Participants performed the task well. On average, they duly switched on most pro-Action trials (*M* = 89.58%, *SD* = 6.03%) and switched rarely in anti-Action trials (*M* = 3.04%, *SD* = 2.14%). Trials with errors of omission and commission were excluded from EEG analysis. Thus, all Informative trials reported correspond to Action trials in the pro-Action condition, and No-Action trials in the anti-Action condition. Participants decided to switch the colour of the screen more often (t(_15_) = 3.31, *p =* 0.004) and sooner (t_(15)_ = 2.46, *p* = 0.026) in the NH (Percentage of Action trials: *M* = 40.54 %, *SD* = 10.89%, Time of action: *M* = 17.50 s, *SD* = 2.20 s), than in the NL condition (Percentage of Action trials: *M* = 34.13%, *SD* = 8.25%; Time of Action: *M* = 18.89 s, *SD* = 1.56 s). However, the average number of evidence items seen before action in the NH condition (*M* = 14.94, *SD* = 1.24) was significantly higher (*t_15_*) = 26.06, *p* < 0.001) than in the NL condition (*M* = 9.19, *SD* = 0.84), suggesting that participants adjusted their decision criterion to the amount of available evidence (see *Figure 2a*). Since the net balance of +Ev and –Ev was the same in those two conditions, this strategic adjustment must have depended on an internal signal. We explore this hypothesis in the computational section below.

**Figure 2.**
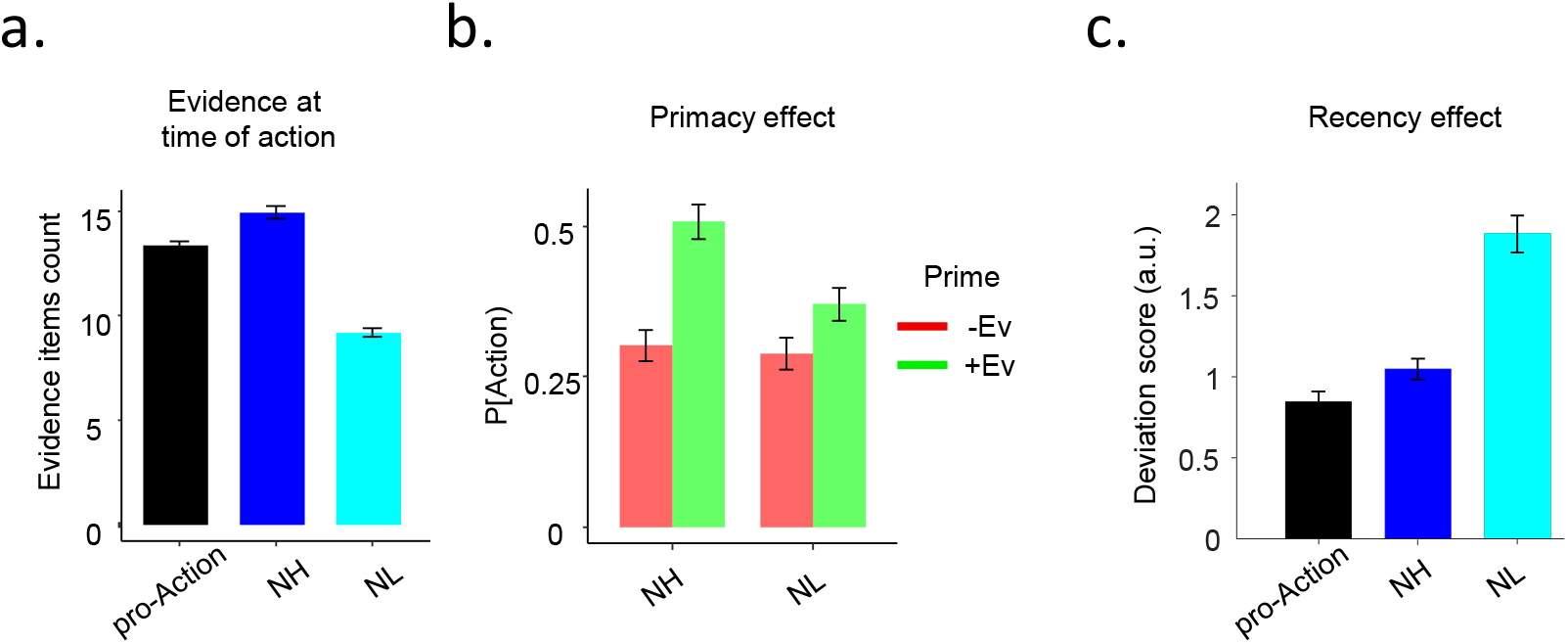
Behavioural results. **a.** Grand-averaged (±SEM) number of evidence items presented at the time of action for all conditions. Although the net balance of +Ev and –Ev was equivalent in the NH and NL conditions, a smaller number of evidence items sufficed to trigger Action in the NL condition. **b.** Grand-averaged probability (±SEM) of Action decisions given the first letter seen on a trial, for NH and NL conditions. In both conditions, participants were more likely to act if the first target letter they saw on a trial was +Ev (e.g. ‘b’ /‘d’ in a pink screen, or ‘p’/’q’ on a blue screen). **c.** Grand-averaged (±SEM) effect of recent evidence on action triggering (see *Figure S1* for full details). Higher values of the Recency Index (RI) indicate a greater dependence of action on recent evidence. In the NL condition, single pieces of evidence items were more likely to be followed by an action than in the other two conditions.

Participants’ subjective reports of their action times significantly preceded the actual movement time by an average of 2.62 s (*SD =* 0.23) across all conditions (t-test against zero: *t*_(15)_ = 11.33, *p* < 0.001). Unsurprisingly, participants were more confident about their decisions in the informative (*M* = 67.55, *SD* = 3.99) than both in the NH (*M* = 47.56, *SD* = 4.35; *t*_(15)_ = 11.90, *p* = < 0.001) and NL (*M* = 48.79*, SD=*4.11; t_(15)_ = 10.54, *p* = < 0.001) conditions, which did not significantly differ between them (t_(15)_ = 1.11, *p* =0.281).

We then investigated how evidence presented early and late in the trial influenced participants’ decisions in our neutral conditions. We found that temporal fluctuations of evidence occurring e rly in the tri l strongly influenced p rticip nts’ decision to ct or not ct (*primacy effect,* see *Figure 2b*). In particular, the first evidence item participants saw in each trial (+Ev/-Ev) biased their Action or No-Action choices in neutral conditions (NH/NL). A main effect of the prime on the probability of deciding to act (*X*^2^_(1)_ = 20.48, *p* < 0.001) showed that participants decided to switch the colour of the screen more often when the first letter they saw on that trial was +Ev (NH: *M* = 50.84%, *SEM* = 2.90%; NL: *M* = 37.10 %, *SEM* = 2.75%) than when it was –Ev (NH: *M* = 31.08%, *SEM* = 2.57%; NL: *M* = 31.21%, SEM = 2.62%) in both the NH and NL conditions. Importantly, the first target stimulus seen on any given trial was independent of the condition and the colour of the screen. A significant interaction between the first letter seen on the trial and the condition indicated that the primacy effect was stronger in the NH condition (*X*^2^ _(1)_ = 5.48, *p* = 0.019).

We further quantified the extent to which the time of actions was influenced by immediately-preceding evidence (*recency effect*). We assumed that, if participants were making a decision completely independently of the environment, the temporal distribution of +Ev and –Ev events prior to action should be uniform distributed, as expected at random. However, if their actions depend on immediate external evidence, the observed distribution just prior to actions should deviate from random. We therefore compared the observed distribution of –Ev (i.e. evidence *against* making a key press) and +Ev (i.e. *for* making a key press) to the expected uniform distribution during the 2.5 s preceding an action (see *Methods* for full details) and obtained a summary measures (*Recency Index, RI*). The greater the RI, the greater the dependency of actions on immediate evidence. In all conditions, participants tended to move shortly after a piece of +Ev (*Figure 2c*; and *Figure S1* for full details). However, the deviation from the expected distribution was greatest in the NL condition (*F*_(2,15)_ = 10.59, *p* < 0.001; see *Supplementary note 1* for full statistical details).

These behavioural results therefore suggested two primarily different mechanisms of symmetry-breaking in the neutral conditions: a confirmation bias driven by early evidence when evidence was abundant (NH), and impulsive reaction-like responses to late evidence when evidence was scarce (NL).

### 2.1.3. The P3 encodes a categorical decision variable

Based on previous literature suggesting that the P3 encodes a decision variable (Twomey et al., 2015), we first investigated the evolution of P3 amplitude in response to sequential, discrete presentations of target letters (i.e. ‘b’,’d’,’p’ and ‘q’). We hypothesized that the evolution of sequential P3 components throughout each trial would exhibit a pattern of build-up associated with participants’ decision-making process. In particular, we hypothesized that the P3 evoked by each stimulus would reflect the accumulated evidence for that hypothesis (+Ev, evidence for Action; –Ev, evidence for No-Action), and thus encode a decision variable driving participants’ choices on a trial-by-trial basis. Since the time of decision in the Noaction trials is unknown, all stimuli up to the end of the trial were included for the sequential P3 analysis. In Action trials, only stimuli up to the time of decision were included.

To study the dynamics of evidence accumulation as encoded in the P3, we extracted the P3 at Pz in response to every instance of relev ant evidence (‘b’, ‘d’, ‘p’ and ‘q’), and we obtained the average amplitude of the whole P3 component (0.3 to 0.8s post stimulus). *Figure 3a* shows the evolution of sequential stimulus-locked P3 amplitudes, as a function of the time within the trial that each target letter appeared. In Action trials, an overall increase in the P3 amplitude is observed from trial onset, peaking at the time of the action. This increase is driven by the increase in the P3 amplitude in response to evidence in favour of the eventual decision to act (+Ev, *Figure 3b*). In No-Action trials, the time of any decision to withhold action is unknown, since participants were not required to produce any response. However, the evolution of the P3 amplitude throughout those trials exhibited a pattern similar to that observed in Action trials. That is, the overall P3 amplitude increased from trial onset, reaching a peak between 10 and 20 s, and was characterised by a greater increase in the P3 amplitudes evoked by evidence supporting the eventual decision (i.e. in this case evidence *against* action, –Ev).

**Figure 3.**
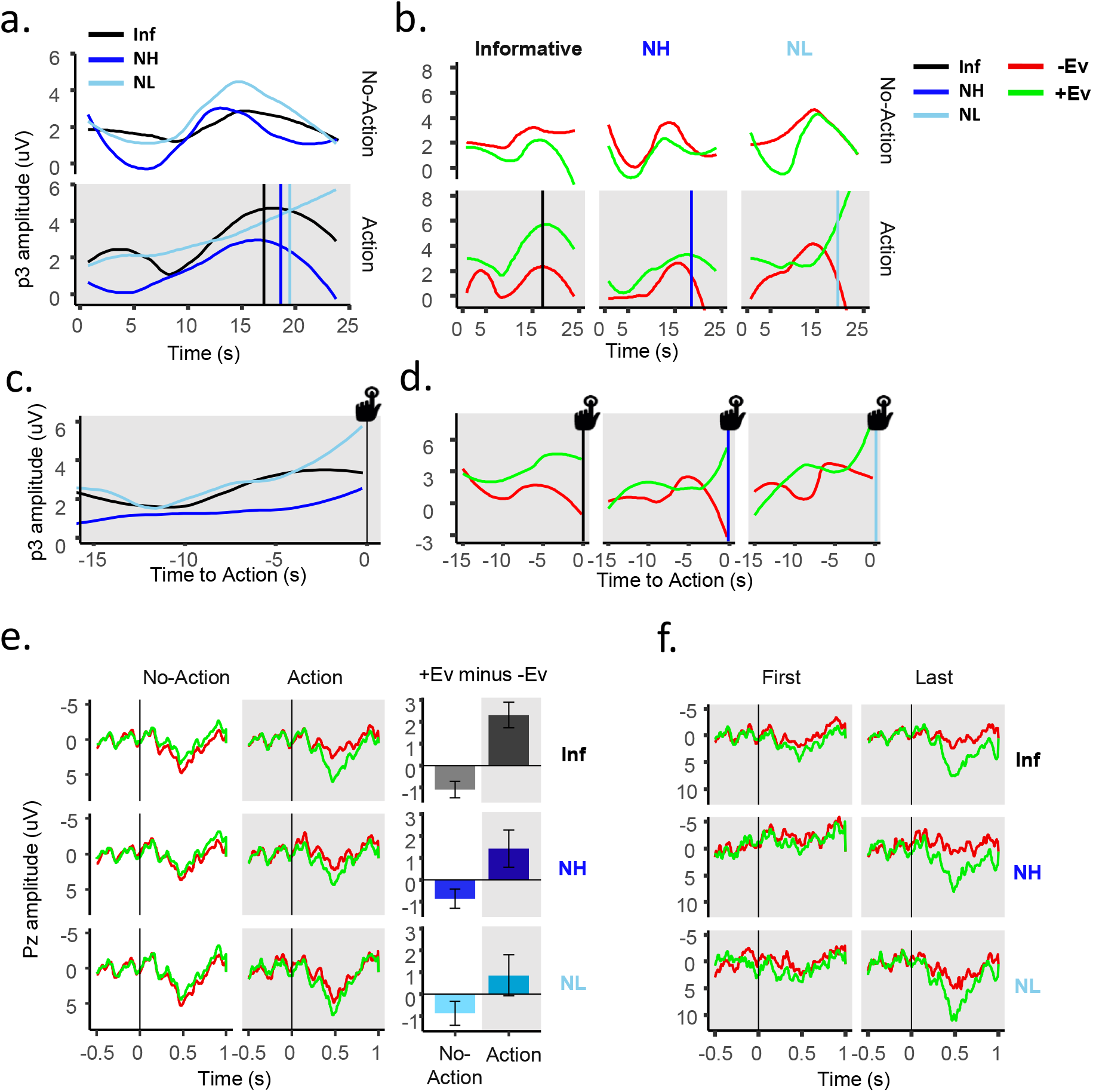
The p3 encodes a categorical decision variable. **a.** Stimulus-locked evolution of p3 amplitude at Pz from trial onset, pooled across participants and types of evidence in Action (*top*) and No-action (*bottom*) trials (loess smoothed). The amplitude of the p3 increased over time in all conditions, although descriptively less in No-action trials. **b.** Evolution of the p3 amplitude at Pz from trial onset, pooled across participants for each type of evidence separately in No-Action (*top*) and Action (*bottom*) trials (loess smoothed). Vertical bars indicate mean time of Action in each condition. **c.** Evolution of p3 amplitude at Pz locked to action in Action trials, pooled across participants and types of evidence. **d.** Evolution of the p3 amplitude at Pz locked to action in Action trials, pooled across participants for each type of evidence separately. Solid lines correspond to the observed data (loess smoothed), and dashed lines to the model prediction. The amplitude of the p3 only increased significantly in response to +Ev. **e.** ERP traces (*left*) show the grand-average p3 in No-Action and Action trials, for +Ev and –Ev. Bar graphs (*right*) show the trial-by-trial averaged difference (+SE) between +Ev and –Ev (+Ev minus –Ev), showing that evidence in favour of the selected option (i.e. +Ev in Action, –Ev in No-Action) evoked significantly higher p3 amplitudes on average, within any given trial. **f**. p3 amplitude in response to the first (*left*) and last (*right*) piece of +Ev and –Ev presented before action in Action trials only. Note: all shaded graphs correspond to trials where an action was executed and include p3 data only up to the time of movement.

To investigate the build-up dynamics of decision-relevant information prior to action, we aligned the P3 data to the time of action on Action trials only (*Figure 3c, d*). We fitted a linear multilevel regression to predict the amplitude of the P3 evoked by each target letter (*bd/pq*) using a linear and a quadratic parameter indicating the time to action (in seconds), the type of letter that evoked it (+Ev/-Ev), participants’ decision (Action/No-Action), the condition (Informative/NH/NL) and all possible interactions as predictors. A significant interaction between the time to action and type of evidence (*X*^2^_(1)_ = 3.98, *p* = 0.045) revealed that the amplitude of the P3 significantly increased in response to +Ev towards the time of action (*X*^2^_(1)_ = 10.20, *p* = 0.001), but not in response to –Ev (*X*^2^_(1)_ = 0.12, *p* = 0.732). To visualise this gradual increase of the P3 in response to +Ev towards the time of action, we plotted the ERPs for the 3 target letters immediately preceding action (*Figure S2*). See *Table S1* for full statistical details, and *Figure S3* for the unsmoothed data used in the analysis.

We further analysed the trial-by-trial P3 amplitudes to investigate whether the selective increase in P3 amplitude would be reflected in the averaged ERPs (*Figure 3e*). We averaged all P3 amplitudes for each trial (i.e. up to the time of action in Action trials, and up to the end of the trial in No-action trials) and fitted a linear mixed model to predict the mean P3 amplitude based on the type of evidence (+Ev/-Ev), the decision (Action/No-Action) and the condition (Informative/NH/NL), ignoring the time at which evidence was presented. We found an interaction between the decision and the type of evidence (*X*^2^_(1)_ = 31.56, *p* < 0.001). In trials in which participants decided to switch the colour of the screen (Action), the amplitude of the P3 was significantly higher (*X*^2^_(1)_ = 18.14, *p* < 0.001) in response to +Ev (*M* = 2.73 μV, *SEM* = 0.43 μV) th an to –Ev (*M* = 0.86 μV*, SEM* = 0.47 μV). In trials in which participants decided to not switch the colour of the screen (No-Action), the effect was in the opposite direction (*X*^2^_(1)_ =12.82, *p* < 0.001). The amplitude of the P3 in response to +Ev (*M* = 1.16 μV*, SEM* = 0.38 μV) was, on average, lower than that in response to –Ev (*M* = 1.96 μV*, SEM* = 0.36 μV). There was no three-way interaction between condition, type of evidence and decision (*X*^2^_(1)_= 3.23, *p* = 0.198), suggesting that the effect was present in both easy (*Informative*) and difficult (*Neutral*, NH & NL) trials (see *Table S2* for full statistical details). This shows that the P3 encoded an internal decision variable, rather than simply the amount of external decision-relevant evidence. Note that given the trial-by-trial variability, the effect is more apparent in the trial-by-trial differences (bar graphs in *Figure 3e, right*) than in the grand-averaged ERPs (*Figure 3e, left*).

Finally, we compared the initial and final amplitudes reached by the P3 in Action trials (*Figure 3f*). If the P3 genuinely encodes a categorical decision variable, accumulator models predict that differences between alternative options should be minimal at the beginning of the decision-making process, and maximal at the time a decision is made. That is, while the P3 responses to the *first* (initial) pieces of +Ev and –Ev presented in a trial should not differ, the P3 responses to the *last* (final) pieces of +Ev and –Ev should reflect the decision participants will make. We fitted a mixed regression to predict the P3 amplitude based on the time within the trial (first/last), the type of evidence (+Ev/-Ev) and the condition (Inf/NH/NL). An interaction between evidence type and time within the trial (*X*^2^_(1)_ = 3.97, *p* = 0.046) revealed that that while at the beginning of the trial the P3 responses to +Ev and – Ev did not significantly differ (*X*^2^_(1)_ = 2.73, *p* = 0.097), the P3 responses to the last piece of +Ev were significantly higher than responses to the last piece of –Ev (*X*^2^_(1)_ = 22.45, *p* < 0.001).

### 2.1.4. The evolution of the categorical decision variable encoded by the P3 is modulated by a context-dependent urgency signal

We next explored whether participants’ behaviour and the dynamic changes in the P3 response could be captured by computational models of evidence accumulation. We modelled the integration of evidence over time using two competing accumulators (Usher & McClelland, 2001).

In our first model (*Constant gain*), the decision parameters were not allowed to vary across conditions and the weight given to incoming evidence (gain) was constant over time. Based on previous literature (Braunlich & Seger, 2016; Cisek et al., 2009; Murphy et al., 2016; Thura & Cisek, 2014), in the second model (*Rising urgency*) we incorporated an growing urgency parameter that dynamically modulated gain. As urgency monotonically increased over time, gain for both evidence accumulators also increased. This time-varying gain means that, as time elapses, less evidence is required to reach a decision. Finally, in the third model (*Context-dependent urgency*), we allowed the urgency parameter, and thus the increase in gain over time, to vary separately for each condition. Condition-specific modulations in an internal urgency signal could potentially account for the different decision thresholds observed in the NL and NH conditions (*Figure 2a*).

Both urgency models performed significantly better than the first model, but did not differ significantly between them. In aggregate, the *Context-dependent Urgency* (total AIC = 6638.40) was numerically better than the *Rising Urgency* model (AIC= 6664.76). Further, the context-dependent urgency model was the best performing one for the majority of participants (n = 12 of 19). This suggests that condition-specific modulations of evidence accumulation occurred, and that they may be explained by a context-dependent urgency signal. Full details of the model specification, comparison and parameter estimation can be found in the Supplementary Material. Below we describe the *Context-dependent Urgency* model in more detail.

The context-dependent urgency model accurately predicted the probabilities of action and the reaction times for all 19 participants (*Figure 4a*). The estimated urgency parameters in this model indicated that the weight given to individual pieces of evidence was greater in conditions where evidence was less frequent (*Figure 4b*). When the rate of evidence was high, the estimated urgency was low (Urgency_NH_ = 0.009). Instead, when the rate of evidence was low, the estimated urgency was higher (Urgency_NL_ = 0.045). This means that while in the NH condition a decision variable evolved towards a threshold mostly driven by the accumulation of abundant external evidence, in the NL condition the decision variable approached the threshold thanks to a strong contribution from an internal urgency signal, which boosted the impact of the scarce available evidence in the accumulation process. Such a modulation can account for the fact that participants acted after having seen significantly fewer evidence items in the NL condition as compared to the NH one (see *Figure 2a*). The inhibition parameter was on average greater than 0 (*M* = 0.19, *SD* = 0.12), t_(18)_ = 6.58, p < .001, and was greater than 0 for 16 out of 19 participants, indicating lateral inhibition between accumulators.

**Figure 4.**
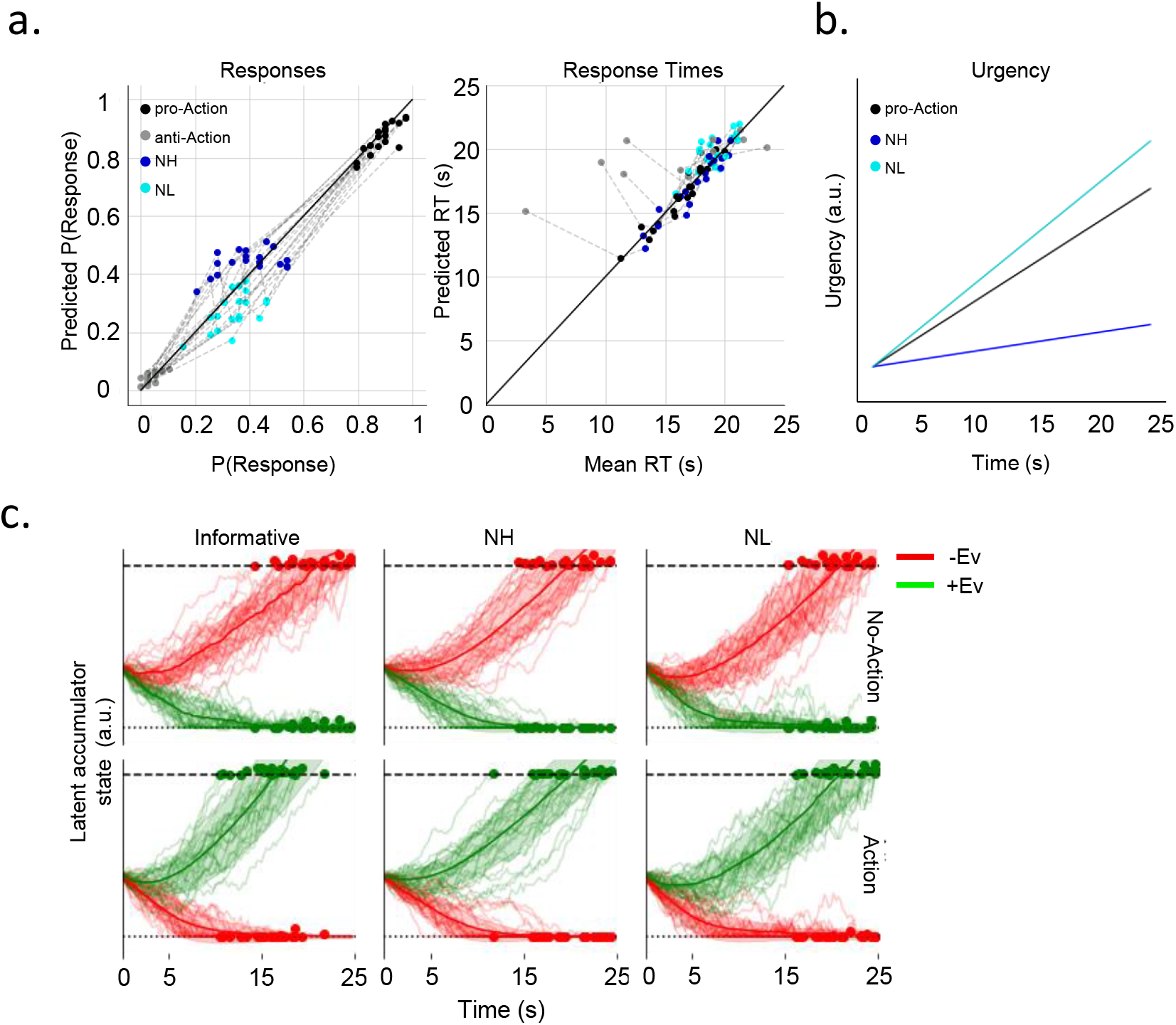
Behavioural predictions and dynamic simulations of the context-dependent urgency model. **a.** Observed and predicted response probabilities (*left*) and response times (*right*) for each condition separately. **b.** Estimated evolution of the urgency signal in arbitrary units (a.u.) for each condition separately. The urgency was estimated to be highest for the NL condition, where evidence was scarce, and lowest for the NH condition, where evidence was abundant. **c.** Simulated accumulator states over time recapitulate our analyses of P3 amplitudes in response to +Ev and –Ev evidence (compare *Figure 2b*). The state of the +Ev accumulator is greater than that of the –Ev accumulator on trials when action occurs (*bottom,* Action), and vice versa when no action takes place (*top,* No-Action). The magnitude of this effect increases as the time advances.

To explore the dynamics of the model, we set each parameter to its average across participants and we simulated 1,000 trials per condition (*Figure 4c*). These simulations broadly recapitulate the P3 evolution during trials (*Figure 2b*). On trials where participants act, the activity of the +Ev accumulator (P3 responses to +Ev stimuli) is stronger than that of the –Ev accumulator (responses to –Ev stimuli). On trials where participants do not act, this effect is reversed. Furthermore, the difference between the two accumulators is weak at the start of each trial and increases over time.

### 2.1.5. Markers of self-paced action in evidence-informed decisions

Finally, we investigated the neural activity preceding action execution to test whether the readiness potential would appear in all, some or no conditions in our task. We found an RP preceding actions in all conditions. We performed pairwise cluster comparisons between all conditions including the two seconds preceding the time of action (−2 to 0). An RP-like shape was visible in all conditions (*Figure 5*), and pairwise cluster comparisons between all conditions found no significant clusters in any of the comparisons (all *p >* 0.05), suggesting that the RPs were not significantly different in amplitude in the different conditions.

**Figure 5.**
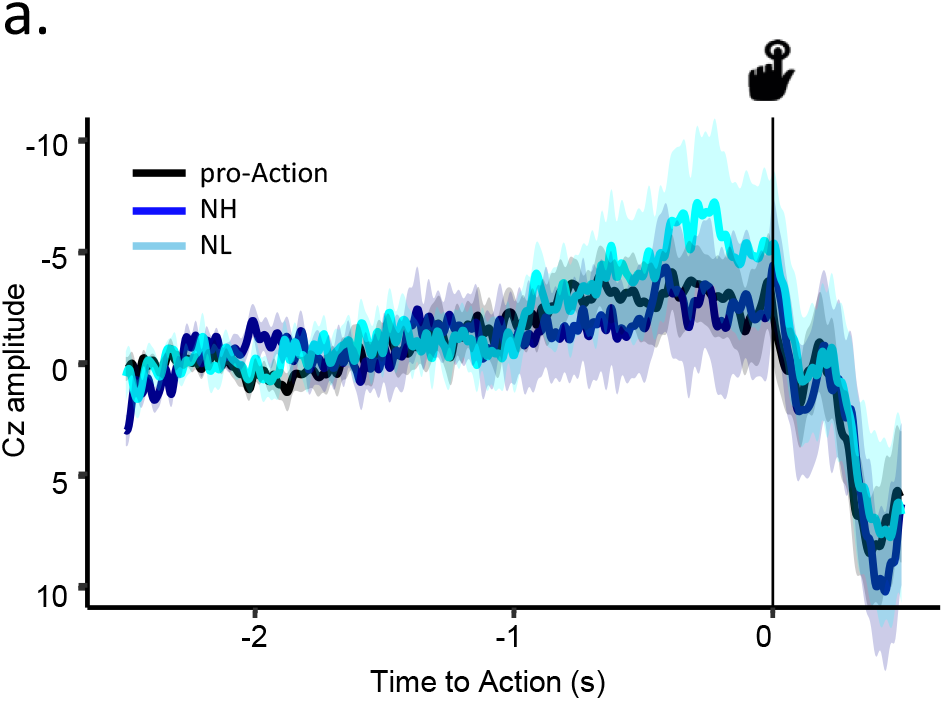
The RP preceding evidence-informed voluntary actions. Grand-averaged EEG activity at Cz preceding actions in all conditions.

## 3. Discussion

The ability to decide when faced with ambiguous information is crucial for functional voluntary behaviour. In this experiment, we developed a paradigm to investigate the neural mechanisms enabling decision and action under uncertainty. In our task, participants watched a stream of discrete evidence items, and had to decide whether to act or not based on the accumulated evidence. However, actions preserved two aspects traditionally present in conventional voluntary action experiments: the fact that decisions might be underdetermined by exogenous evidence, and the self-paced nature of the movement. Our goal was to investigate the neural mechanisms that allow symmetry-breaking at the sensory and motor levels. We focussed own the P3 component as a marker of evidence accumulation processes and the RP as a marker of self-paced motor preparation.

### The P3 encodes an internal categorical decision variable

Our use of sequential category-specific stimuli rather than continuous stimulation (cf. O’Connell et al. 2012) allowed us to investigate the P3 neural response evoked by each item as a putative neural correlate of accumulating evidence for specific choices. To analyse the decision-making process throughout the long trial durations, we developed a novel sequential ERP analysis method. Rather than investigating the peak latency and slope of an average ERP trace, (Kutas et al. 1977, Twomey et al. 2015), we analysed how the P3 component evoked by successive stimuli evolved over time during the decision-making processes, for each trial individually. We show that sequential P3 amplitudes show build-to-threshold dynamics, reaching peak amplitude at the time of action (Action trials), and at the presumed time of decision in No-Action trials (*Figure 3a*). By aligning the evolution of the discrete P3 responses to the time of action (in Action trials), we showed that the P3 evoked by sequential stimuli selectively increased in response to +Ev towards the time of the action (*Figure 3*). The P3 amplitude initially increased for both types of evidence, but then diverged. Interestingly, these dynamics are similar to single-cell recordings in monkey’s lateral intra-parietal (LIP) area during a decision-making task (Churchland et al., 2008; Huk & Shadlen, 2005; Roitman & Shadlen, 2002). The same P3 dynamics throughout single trials and average ERPs were visible in *all* conditions, including neutral trials which contained no net evidence. Since the P3 evolution nevertheless reflected participants’ choices, it seems to track the state of an internal decision variable rather than objective evidence.

Participants’ choices were also reflected on the average of P3s evoked b y *all* stimuli within a trial (*Figure 3e*). In trials where participants decided to act, the amplitude of the P3 in response to evidence favouring Action (+Ev) was significantly greater than the P3 amplitude in response to evidence favouring No-Action (–Ev). In No-action trials, this pattern was reversed. The P3 thus encodes a decision variable even in trials in which no motor response is required (i.e. No-action trials), in agreement with previous literature (O’Connell et al. 2012). The fact that the P3 amplitude reached a peak on average well before the end of these No-Action trials further suggests that participants did not continue to accumulate evidence until the end of the letter stream, but rather made a covert decision to *not* move before hand and then stopped accumulating evidence. This is in agreement with previous literature showing that once a decision bound is reached, further evidence is not further processed for perceptual decisions (Balsdon et al., 2020; Kiani et al., 2008). The response-independence of the P3 suggests that it carries information about an intermediate stage between sensory evidence and motor action. Further, the fact that the P3 response to an individual stimulus depends on the decisional category to which the stimulus belongs is consistent with findings that single-neuron firing patterns specifically represent distinct decision alternatives in the monkey parietal cortex (Fitzgerald et al., 2011; Freedman & Assad, 2006; Swaminathan & Freedman, 2012). To our knowledge, this is the first demonstration that the P3 component may reflect such categorical processing in humans.

### The categorical decision variable encoded by the P3 is modulated by a context-dependent urgency signal

Which signals and neural mechanisms drive the evolution of the decision variable that the P3 encodes? Our best-fitting model showed that participants’ behaviour was best explained by a competing accumulation process with a condition-specific gain adjustment, mediated by a growing urgency signal. In particular, we showed that evidence processing was tuned to the availability of relevant information in each condition (see *Figure 4b*).

Urgency is often conceived as an evidence-independent, time-growing signal that facilitates making a decision in various contexts by modulating the amount of evidence required for a decision as time elapses (Braunlich & Seger, 2016; Churchland et al., 2008; Ditterich, 2006; Thura, 2020), and it has been shown to depend on the basal ganglia (Bogacz et al., 2010; Thura & Cisek, 2017). Here, we followed studies showing a multiplicative interaction between evidence accumulation processes and urgency signals (Cisek et al., 2009; Ditterich, 2006; Thura & Cisek, 2014). In our task, conditions differed not only in the balance, but also in the amount of evidence available. Thus, we further hypothesized that the different conditions effectively constitute different contexts where the rate at which urgency grows, may depend on evidence availability. Indeed, our best-fitting model showed that when evidence was scarce (NL), the urgency signal was estimated to be highest (*Figure 4b*). Since we postulated that urgency would multiplicatively modulate the gain of the evidence accumulation process, this could be seen as a strategy for reaching a decision when evidence is rare: by increasing the gain in such contexts, each new piece of evidence makes an enhanced contribution to the decision variable. Thus, fewer evidence items will therefore suffice to reach a decision threshold (see *Figure 2a*). In turn, reducing gain in contexts where evidence is abundant allows the accumulation of more pieces of evidence, thus preventing potentially hasty and erroneous decisions.

The simulations based on the population averages of our context-sensitive urgency model further illustrate that the model not only fits behavioural, but also neural data. The evolution of the P3 over trial time broadly captures the latent state of the accumulators. The model predicted a selective increase in amplitude of the P3 response to +Ev (*Figure 4c*), recapitulating the observed evolution of the category-specific P3s (*Figure 2b*). However, the P3 cannot track *only* the mere amount of accumulated evidence items, since the amplitude of the P3 evoked by, say, the third evidence item in the NH condition is substantially lower than that evoked by the same item in the NL condition (see *Figure S4*). In turn, the P3 seems to reflect not only the growing urgency signal, but *also* evidence accumulation. If the P3 reflected urgency only, we would expect the P3 traces to be modulated by the time at which they were evoked, but not by which type of evidence evoked it. In fact, the evolution of the P3 evoked by category-specific stimuli diverged according to participnts’ choices.

In sum, we believe that the amplitude of the P3 evoked by any given stimulus can be explained as a function of 1) the current state of an internal evidence accumulator, which is in turn modulated by 2) a context-dependent urgency signal growing over time. *Figure 6* illustrates a schematic of the cognitive mechanisms underlying the decision-making process that we believe the P3 tracks. We suggest that the P3 reflects the activity of category-specific neural populations encoding incoming evidence. Further, the availability of evidence modulates the rate at which an internal urgency signal grows, which in turn modulates the gain (*g*) of the evidence accumulators. As a decision-making process about the distribution underlying the stimuli evolves (i.e. are *bd* more frequent than *pq*?), the closer the accumulated evidence (∑) moves towards threshold. It remains possible that additive urgency modulations were also in place, as additive and multiplicative effects are not necessarily mutually exclusive (Murphy et al. 2016). The goal of our modelling was only to show that multiplicative urgency modulations provide a plausible account of the observed data.

**Figure 6.**
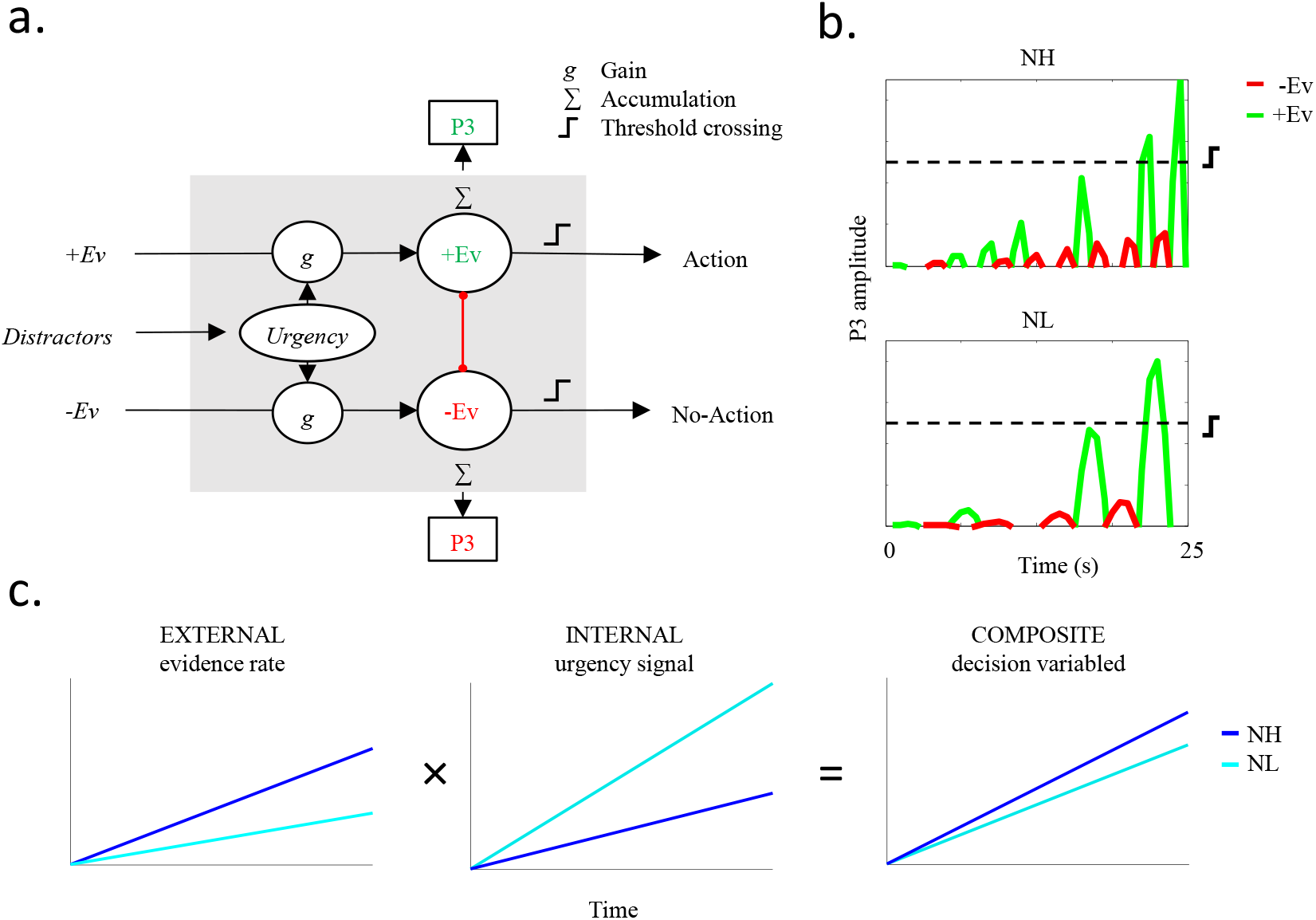
Conceptual model of the cognitive mechanisms underlying the P3. **a.** Suggested computational mechanisms driving evidence accumulation in our task. Evidence is processed sequentially, and each stimulus is processed with a certain gain (*g*) in a category-specific accumulator. The accumulators for competing choices are mutually inhibitory. The gain in turn depends on a time-varying urgency signal. The rate at which urgency, and hence gain, increases over time is a function of the availability of relevant evidence. The more evidence is available, the smaller the steepness of the urgency signal, allowing the accumulation of more evidence items before reaching a threshold. Once one of the accumulators reaches a threshold a decision is made. **b.** Schematic representation of the evoked P3 responses in a condition where evidence is abundant (NH, *top*) or rare (NL, *bottom*). The evolution along the x axis illustrates the P3 amplitudes over time, which are a function of a competitive evidence external accumulation process modulated by a growing internal urgency signal. In turn, this enables crossing the threshold after fewer evidence items have been presented. The fact that the P3 amplitudes evoked by +Ev and –Ev are different despite the absolute numbers of evidence being the same shows the effect of lateral inhibition. **c.** Schematic of the multiplicative interaction between external evidence and context-dependent internal urgency signal to produce a composite decision variable. When evidence is abundant (NH), the urgency signal grows slowly and the evolution of the decision variable is mostly driven by external evidence. When evidence is scarce (NL), the urgency signal grows faster and its contribution to the evolution of the decision variable is greater.

### Symmetry-breaking mechanisms

Our behavioural results in the neutral conditions suggested that two seemingly different mechanisms might account for symmetry-breaking. On the one hand, asymmetries at the start of an evidence accumulation process may determine the course of subsequent evidence accumulation and hence the final decision (*primacy effect*). On the other hand, a single stimulus item may suffice to trigger action (*recency effect*) and resulting in more impulsive, reaction-like actions. Can the neural mechanisms of decision making described above account for these behavioural patterns in the neutral conditions of our task?

In neutral trials, participants were more likely to decide to act if the first target letter that they saw provided evidence for action, rather than inaction - irrespective of what evidence they saw subsequently (*Figure 2b*). Competing accumulator models like the ones we explored one predict this kind of primacy effect (Tsetsos et al., 2012). Small differences at the beginning of an accumulation process are amplified over time thanks to the mutually inhibiting connections between competing accumulators. This symmetry-breaking mechanism is thus like a neural confirmation bias: in the absence of strong countermanding evidence, decisions outcomes tend to follow early hypothesis. In our study, these asymmetries depended on the time at which external stimuli were presented. However, early brain activity has been reported to contain information that enables decoding free decisions in humans (Soon et al., 2008), and it has been suggested that free decisions depend on accumulation processes similar to those operating in perceptual ones (Bode et al., 2013). Thus, early asymmetries of either endogenous or exogenous origins can effectively bias accumulation processes and thus act as symmetry-breakers.

In turn, we found that actions typically shortly followed the presentation of a +Ev item (*Figure S1*). That is, the time of actions was dependent on immediate evidence to a certain extent in all conditions. This is to be expected, since participants decisions were not independent of the evidence presented on screen. However, the influence of immediate evidence on the time of actions was strongest in the NL condition, where external evidence was particularly sparse (*Figure 2c* and *S1*). Such effect can be explained by the condition-specific modulations of urgency that we modelled. The increased urgency estimated for the NL condition means that the impact of any individual piece of evidence on the decision variable is comparatively high. Therefore, the probability that any given evidence item will make the decision variable cross the threshold is highest in that condition, accounting for the recency effects.

In sum, our behavioural results can be accounted for by our suggested computational model: when evidence is abundant decision variables evolve up to threshold driven mostly by evidence itself, even when this evidence is neutral. In turn, when evidence is scarce decision variables raise to threshold driven mostly by an internal urgency signal that boosts evidence processing. This leads to the paradoxical conclusion that when people base their decisions on less external evidence they are also more reactive, and therefore less ‘free from immediacy’ (Gold & Shadlen, 2003)

### The RP can be found preceding evidence-informed actions

Finally, we investigated the neural precursors of action initiation. Classic interpretations of RP suggest that it reflects endogenous neural activity leading to action. Research has typically measured the RP in experiments where the participant acts as and when they wish, without any imperative stimulus or explicit sensory evidence to trigger the decision to act (e.g. Libet et al., 1983). In an attempt to develop a more naturalistic paradigm, our task was designed as a hybrid between classic spontaneous action paradigms and cued-reaction tasks: participants had no strong time pressure to execute actions, and although their actions were informed by the evidence, they were not triggered by any specific individual stimulus.

Readiness potentials were visible preceding actions in all conditions, and the amplitudes did not significantly differ across conditions (*Figure 5*). The fact that an RP was also visible preceding actions in the pro-Action condition, where actions were deliberate and strongly determined by the environment is not compatible with the idea that stochastic accumulation of internal noise that the RP reflects (Schurger et al. 2012) is present *only* in contexts where actions are arbitrary (Maoz et al. 2019).

### Conclusion

In sum, the present study investigated the neural mechanisms that enable symmetry breaking and consequent action during decision under uncertainty. Our results expand the previous knowledge regarding neural correlates of an internal decision variable in several ways. First we show that the P3 evoked by discrete evidence items tracks not only decision time, but also categorical choice. Second, the P3 evoked by sequential pieces of evidence reflects the evolution of a decision, capturing the current accumulated balance of positive and negative evidence. Third, the decision variable captured by the P3 is modulated by a context-dependent urgency signal that depends on the rate of evidence. Further, a combination of competitive evidence accumulation and context-sensitive urgency can account for two seemingly different behavioural symmetry-breaking patterns: a confirmation bias mechanism driven by early evidence (*primacy effect*) and a late reaction-like impulsivity (*recency effects*). Finally, we show that the RP can be found in evidence-driven self-paced actions, validating its role as a marker of voluntary action even in externally-driven actions.

## 4. Methods

### Participants

All participants were recruited from the ICN Subject Database. All participants were healthy, right-handed, young adults with normal or corrected to normal vision, no known disabilities and no history of neurological or psychological disorder. The study was approved by the UCL Research Ethics Committee and written informed consent was obtained from all participants before beginning the experiment. Subjects were paid £7.50 per hour, plus a performance-dependent reward (see below).

Nineteen subjects were initially invited to a single EEG session. Three participants were excluded from EEG analysis due to excessive EEG noise in the electrodes of interest (Cz & Pz). Eventually, 16 participants (13 female) were included in the EEG analysis (*M* age = 22.66, *SD* = 3.19; range: 19-30 years). All participants were included for the computational modelling.

### Stimuli & experimental design

### Procedure

Participants sat in a quiet room and were presented visual stimuli on a computer monitor. The instructions for the task were first displayed on the computer screen and then verbally repeated by the experimenter before the beginning of the experiment. Before the experiment, participants performed a practice version of the task (5 trials) to get familiar with the experiment.

### Stimuli

A continuous stream of pseudo-randomised letters was presented at a 4Hz rate (duration of each letter 250 ms) on either a grey, pink or blue background (see below). Every trial lasted 25 s. The experiment was programmed in Matlab R2015a and Psychophysics Toolbox v3 (Brainard, 1997). Participants made actions by pressing either the ‘j’ (pink) or the ‘f (blue) key of a standard computer keyboard with their left or right index finger respectively and answered the post-trial questions (see below) on a visual analog scale (VAS) by sliding a scroll-bar with a standard computer mouse.

All letters were lower-case black consonants presented without any blank interval between consecutive letters. There were two sets of task-relevant (target) letters, to which a task-relevant colour was assigned (in parenthesis): ‘b’ and ‘d’ (*bd,* blue), and ‘p’ and ‘q’ (*pq,* pink). The precaution was taken not to include letters ‘h’, ‘j’, ‘k’, ‘l’, ‘t’ in the letter stream to ensure that morphological similarities between these and the target letters would not confound the results.

### Reward

In both Informative conditions, participants were rewarded with +1 penny for correct decisions (i.e. acting in the pro-Action trials and not acting in the anti-Action trials) and penalised for incorrect decisions (i.e. acting in the anti-Action trials and not acting in the pro-Action trials) by deducting 1 penny. In the neutral conditions, there were no correct or incorrect decisions. Hence, participants were assigned +1 penny reward or no reward at random on every trial. Participants were informed about the accumulated reward on the breaks between blocks.

### EEG recording

EEG was recorded from 26 scalp sites (Fz, FCz, Cz, CPz, Pz, POz, FC1, FC2, C1, C2, CP1, CP2, F3, F4, F7, F8, C3, C4, CP5, CP6, FC5, FC6, P3, P4, O1, O2) using active electrodes (g.LADYbird) fixed to an EEG cap (g.GAMMAcap) according to the extended international 10/20 system. EEG data were acquired using a g.GAMMAbox and g.USBamp with a sampling frequency of 256 Hz. Signal was recorded using g.Recorder (G.tec, medical engineering GmbH, Austria). All electrodes were online referenced to the right ear lobe. Vertical and horizontal electroocular activity was recorded from electrodes above and below the right eye and on the outer canthi of both eyes.

### Behavioural data analysis

### Perceived time of switch decision

After trials where a decision to switch the colour of the screen was made and therefore participants acted (in Action trials), participants were required to provide an estimate of the time at which they decided to switch. In order to calculate the bias in their estimation with respect to the actual time of the decision to switch we first linearly converted the 100-point VAS on which participants provided the response dividing it by 4, thus providing a 25-point scale equivalent to the 25 s duration of the trial. We calculated the difference between the actual time and the perceived time of action for each individual trial (bias), averaged it across all Action trials for each participant and run a paired samples t-test against zero.

### Primacy effect

To investigate the extent to which decisions in highly ambiguous situations are driven by external evidence, we investigated whether the first piece of evidence participants saw in each trial biased their Action or No-Action choices in neutral conditions. We ran a multilevel logistic regression using the lme4 package (Bates et al., 2015) to predict the probability of Action based on the first piece of evidence (+Ev/-Ev) on each single trial, the condition (NH/NL) and the interaction between both categorical variables. We included a random intercept to account for the between subject variability.

### Recency effect

We developed a measure to estimate the extent to which participant’s actions were dependent on immediate external evidence. We assumed that, if participants were making a decision completely independently of the environment, the distribution of +Ev and –Ev preceding an action should be uniformly distributed, as expected at random. However, if their actions are driven to some extent by immediate external evidence, the observed distribution just prior to actions should deviate from the one expected at random. To measure the magnitude of this deviation, we compared the observed distribution of –Ev (i.e. evidence *against* making a key press) and +Ev (i.e. *for* making a key press) to the expected uniform distribution during the 2.5 s preceding an action, divided in 250 ms time bins. For this comparison, we calculated a *Deviation Score* (DS) for +Ev and –Ev separately (DS_(+Ev)_ and DS_(–Ev)_). For each participant and condition, we subtracted the expected number of ±Ev (±Ev _e_ = probability of any given letter being ±Ev × number of Action trials) from the observed number of (±Ev_o_) in each time bin. We then divided the result by the expected number of targets to normalise the score (Generic formula: DS = (±Ev_o_ - ±Ev_e_) / ±Ev_e_). Positive values in the deviation score indicate that there were more targets than expected, while negative values indicate that there were less targets than expected. Finally, we combined these two deviation scores in a single measure: the *Recency Index (RI*). The Recency Index was calculated by subtracting the DS_(–Ev)_ from the DS_(+Ev)_ (Generic formula: RI = DS_(+Ev)_ - DS_(–Ev)_). The RI is thus a measure of how much the distribution of both +Ev and –Ev combined deviates from the one expected at random. The greater the RI, the greater the dependency of actions on immediate evidence. For statistical analysis, we averaged the RI in each time bin for the whole 2.5 s epoch preceding actions and a repeated measures ANOVA and post-hoc pairwise comparisons for each condition pair.

### Statistical analysis

Statistical analysis of the recency effect described above was run on Matlab R2014b. All other analysis of behavioural data were performed in R using mixed-models regression with the *lme4* (Bates et al., 2015) and *blme* packages. We fit models to compare the percentage of actions and the time of action, and all models included a random intercept to account for the between subject variability.

### EEG analysis

### Preprocessing

EEG data were processed using Matlab R2014b (MathWorks), EEGlab (Delorme & Makeig, 2004) and the Berlin Brain Computer Interface toolbox (Blankertz et al., 2016). First, scalp and eye electrodes were re-referenced to the average of two mastoid electrodes. Continuous EEG and EOG data were low-pass filtered under 30Hz using a zero-phase 8th order Butterworth filter. Second, EEG signals were epoched. For P3 analysis, EEG signal was locked from −0.5 b efore to 1 s after the presentation of relevant letters (‘p’, ‘q’, ‘b’ and ‘d’). For RP analysis, EEG signal was locked from −2.5 s to 0.5 s after the presentation of the letter following a keypress. Next, baseline correction was performed using the 500 ms at the beginning of the epoch [−0.5 to 0 s in P3 epochs, or −2.5 s to −2 s in RP epochs]. Ocular movements were identified using the *runica* algorithm in Matlab and removed from the epoched signal. Finally, artefact rejection was performed by removing all epochs with >200 μV fluctuations from baseline in the preselected channels of interest (CZ and PZ).

### Sequential P3 analysis

To study the dynamics of evidence accumulation as encoded in the P3, we extracted the P3 at Pz in response to every instance of relevant evidence (‘b’, ‘d’, ‘p’ and ‘q’), and we obtained the average amplitude of the whole duration of the component [0.3 to 0.8s post stimulus], as determined by visual inspection of the grand-averaged traces. We used these raw values for the dynamic analysis of the evolution of the P3 amplitude over time in a stimulus-locked (i.e. in absolute time from trial onset) and action-locked manner (i.e. in relative time to execution of action). That is, we investigated how the amplitude of the single stimulus-evoked P3 changed over sequential stimuli presentations throughout the whole trial duration.

For single-trial average ERP analysis, we averaged the P3 components in response to +Ev and –Ev separately for each single trial, and we used these values for statistical inference. We ran a multilevel linear regression using the *lme4* package (Bates et al. 2015). We fit a mixed model to test whether the single-trial average amplitude of the P3 significantly varied between different kinds of evidence (+Ev/-Ev), conditions (Informative/NH/NL) and Action decisions (Action/No-Action).

For initial and final amplitude P3 analysis, we investigated whether the amplitudes of the *first* and *last* pieces of evidence presented from trial onset until action execution or until trial end (in Action and No-Action trials respectively) varied between different kinds of evidence (+Ev/-Ev), conditions (Informative/NH/NL) and Action decisions (Action/No-Action).

In all models, a random intercept was included for each subject and individual trial to control for trial-by-trial and individual variability.

### RP analysis

Statistical tests on averaged EEG data were run using FieldTrip toolbox (Oostenveld et al., 2011) cluster-based permutation analysis (Maris & Oostenveld, 2007). The main contrast of interest was the comparison between neural activity preceding actions between conditions (pro-Action, NH, NL). One electrode over the medial frontal areas (Cz) was preselected for analysis. The cluster-based tests were performed on the individual participant averages using the following parameters: two-tailed dependent samples t-test, time interval = [-2 0 s relative to action], number of draws from the permutation distribution = 10000. No correction for multiple comparisons was performed.

### Computational modelling

### Model description

We modelled the integration of evidence over time using two competing accumulators (Usher & McClelland, 2001). Such a model is a compromise between single-accumulator drift diffusion models (Ratcliff et al., 2016), which track the balance of evidence and therefore cannot account for differences between the Neutral-High and Neutral-Low conditions, and race models (Brown & Heathcote, 2008) which track the evidence in favour of each response but disregard the balance of evidence (Bogacz et al., 2006).

We modelled the state of two accumulators, x^1^ and x^2^ which integrated positive (pro-Action) and negative (anti-action) evidence respectively. The state of the accumulators was updated at every time step, **dt = 0.25**, using the equations:

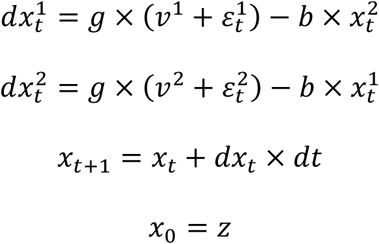

where **g** is a gain parameter, **v^1^** and **v^1^** are the inputs to the positive and negative accumulators respectively, **ε** is independent Gaussian noise with variance **σ**^2^, **b** is a lateral inhibition parameter, and **z** is a starting point parameter. Each condition was simulated separately, with values of **v^1^** and **v^2^** set to match the probability of positive and negative evidence in each condition. These values are therefore not free parameters. We simulated 100 time steps (25 seconds) for each trial. Simulated responses (Action or No-Action) were produced by whichever accumulator was first to reach a criterion of **1**, and simulated response times corresponded to the time at which the criterion was reached. Parameters were estimated separately for each participant.

In the c*onstant gain* model, all free parameters were kept constant across conditions (number of free parameters k = 4, parameter vector **θ** = **[g, b, σ^2^, z]**). In the *rising urgency* model, gain was set to **g_α_** at the start of each trial and increased by **gβ** with each time step (k = 5, **θ** = **[g_α_, gβ, b, σ^2^, z]**). In the *context-dependent urgency* model, **gβ** was estimated separately for each condition, allowing urgency to increase at different rates (k = 8, **θ**^α^ = **[g_α_^Pro-Action^, g_β_^Neutral-High^, g_β_^Neutral-Low^, g_β_^Anti-Action^, g_β_, b, σ^2^, z]**).

### Model fitting

We fit models to each participant’s behavioural data by maximum-likelihood. For a given vector of parameters **θ**, we simulated 500 trials in each condition using these parameters and the appropriate values of **v**^1^ and **v**^2^.

We used the proportion of simulated trials where an action occurred as an estimate of the likelihood of action given these parameters, **P(Act|θ)**. We estimated the response time probability density function, **P(RT = t|Act, θ),** by fitting an Epikanhov kernel density estimate (Turner & Sederberg, 2014) to the distribution of simulated response times on simulated trials where actions did occur. The log-likelihood on each trial was therefore **loglik = log(P(Act|θ)) + log(P(RT = t|Act, θ))** on trials where the participant acted, and **log(1 - P(Act|θ))** on trials where they did not. The total log-likelihood for each participant’s data, **LL = Log(P(Data|θ))** was calculated by summing the log-likelihood of each trial, across conditions. We used differential-evolution optimisation, implemented in the scipy package for python (Virtanen et al., 2020), to maximise total log-likelihood for each participant.

### Model comparison

To compare models, we calculate Akaike Information Criterion (AIC) for each participant, **AIC = - 2×LL + 2×k**, where **k** is the number of parameters per model. Lower AIC values indicate greater support for the model. The differences in the AIC, **δ(AIC)**, were calculated as the difference between the AIC of the model being assessed and the AIC of the best model. These were used to calculate AIC weights, **wi(AIC) = exp{δj(AIC)} / Σ_i_^3^ exp{δj(AIC)}** (Burnham & Anderson, 2001), indicating the probability that each model is the best of the three considered.

**Table 1.**
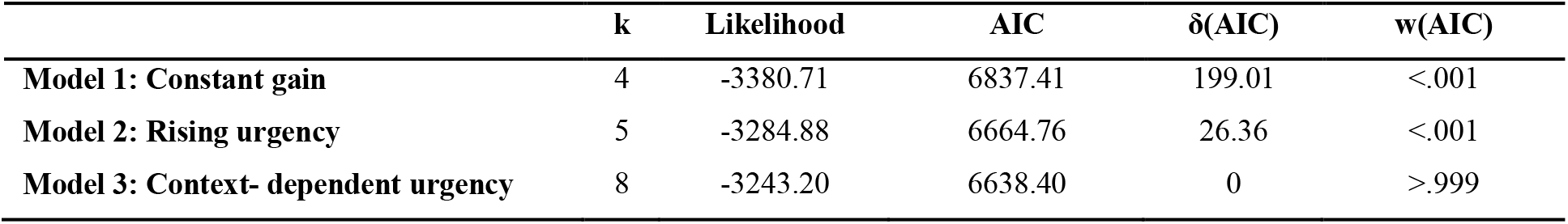

On aggregate Model 3 outperformed the alternatives by a considerable margin, w(AIC) > .999. However, there was some heterogeneity across participants, as Model 3 was superior for 12 participants, Model 2 was superior for 5 participants, and Model 1 was superior for the remaining 2 participants.

## Acknowledgements

This work was partly supported by a joint grant from the John Templeton Foundation and the Fetzer Institute (to PH). The opinions expressed in this publication are those of the author(s) and do not necessarily reflect the views of the John Templeton Foundation or the Fetzer Institute. Furthermore, this work was supported by European Research Council Advanced Grant (project no. 323943), HUMVOL, awarded to PH.

## Authors’ contributions

EPP and PH conceptualized and designed the task. EPP and YA conducted the experiment. EPP and ET analysed the data. EPP, ET, and PH wrote the manuscript.

## 5. Supplementary material

### Supplementary Note 1

To investigate recency effects, we analysed the distribution of +Ev and –Ev presented in the 2.5 s preceding actions (*Figure 3a, top*). We calculated a *Deviation Score* (DS) for +Ev and –Ev separately to measure how much these distributions differed from the one expected if actions were made at random, completely ignoring external evidence (*Figure 3a, middle*). Positive DS values indicate that *more +*Ev or –Ev letters were observed shortly preceding an action, whereas negative values indicate that *fewer* letters than expected at random were observed. On average, there were more were more +Ev and fewer –Ev than expected at random in all conditions – especially during the last second before action execution. Participants were more likely to decide to switch the colour of the screen shortly after a +Ev letter was presented on the screen and less likely to do so when –Ev was presented. We then averaged the two DS for +Ev and –Ev over time and combined them into a Recency Index (RI), measuring how much both distributions combined differed from the expected ones (*Figure 3a, bottom*).

We found a significant effect of condition on the RI (*F(_2,15)_* = 17.52, *p* < 0.001), indicating that actions in the NL (*M_RI_* = 1.88, *SD* = 0.91) condition were more strongly driven by immediate evidence than those in the NH (*M_RI_* = 1.04, *SD* = 0.53; *t*_(15)_ = 6.17, *p* < 0.001) and the Informative pro-Action (*M_RI_* = 0.845, *SD* = 0.50; *t*_(15)_ = 4.46, *p* < 0.001) conditions, which did not differ between them (*t*_(15)_ = 1.14, *p* = 0.269).

**Figure S1.**
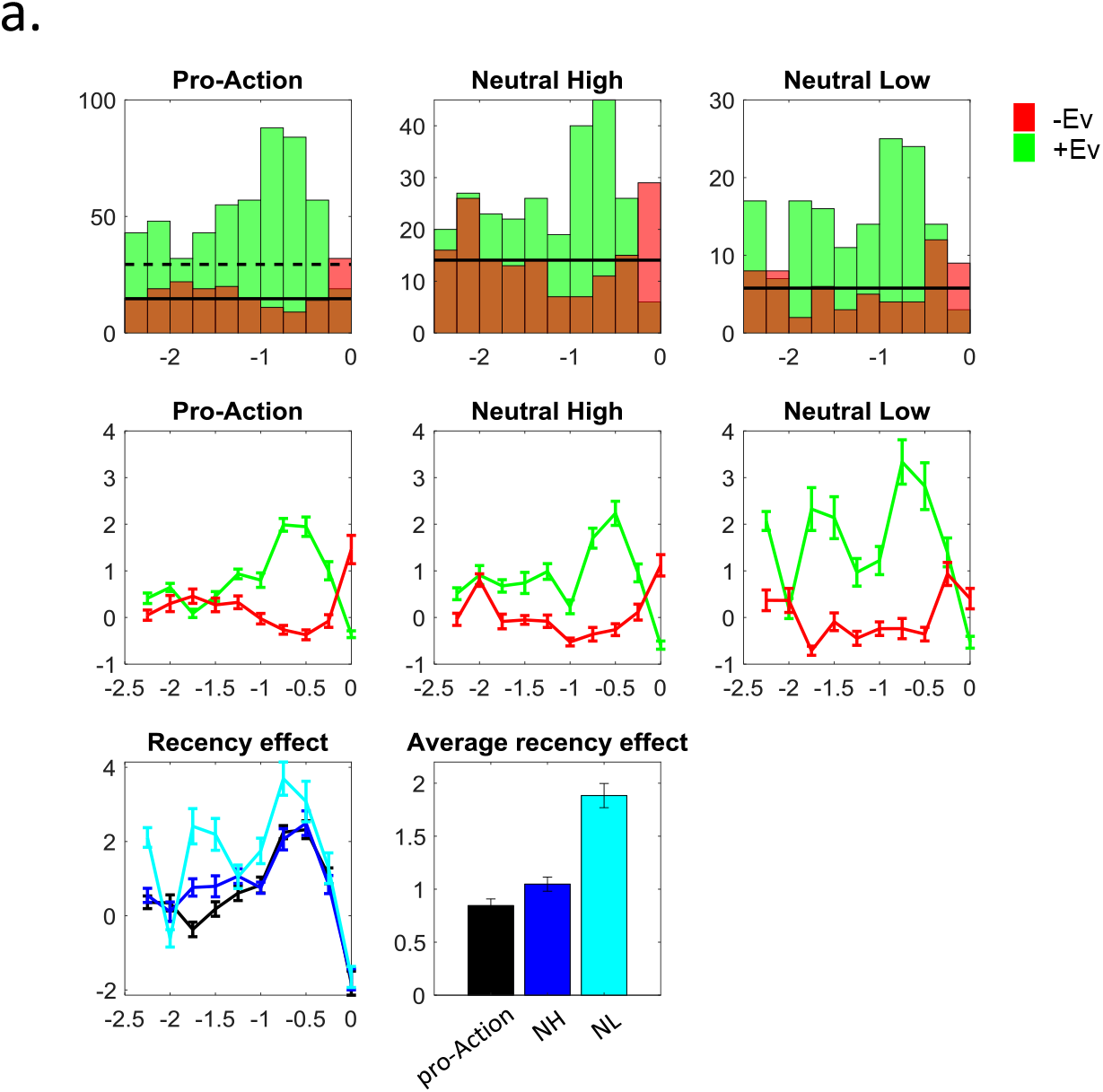
Recency effect. Bar graphs (*top*) show the pooled distribution of +Ev (*green*) and –Ev (*red*) locked to the time of action (0 in the x axis). Horizontal lines indicate the expected probability of each type of evidence. Line graphs (*middle*) show the grand-averaged (±SEM) deviation score (DS), which corresponds to the difference between the observed and the number of +Ev and –Ev letters expected at random during the 2.5 seconds preceding actions for each letter position. Grand averaged (± SEM) Recency Index (RI) over time for the Pro-Action (black), Neutral High (NH, blue) and the Neutral Low (NL, cyan) conditions (*bottom*).

**Figure S2.**
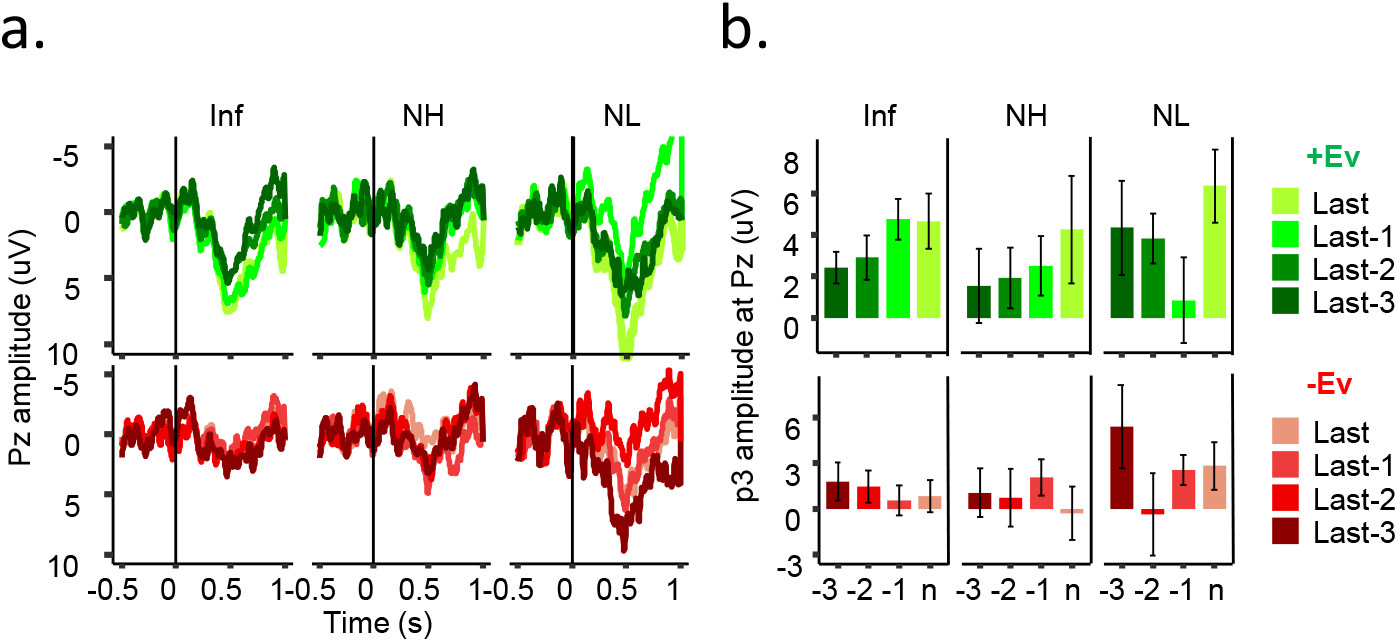
Grand-averaged ERP traces (***a***) and peak p3 amplitudes (***b***) for the last four instances of +Ev (*green*) and –Ev (*red*) presented before actions. Darker colours illustrate earlier events, lighter colurs depict stimuli presented closer to the time of action. Bar graphs represent the grand-average between 300 and 800 ms.

**Figure S3.**
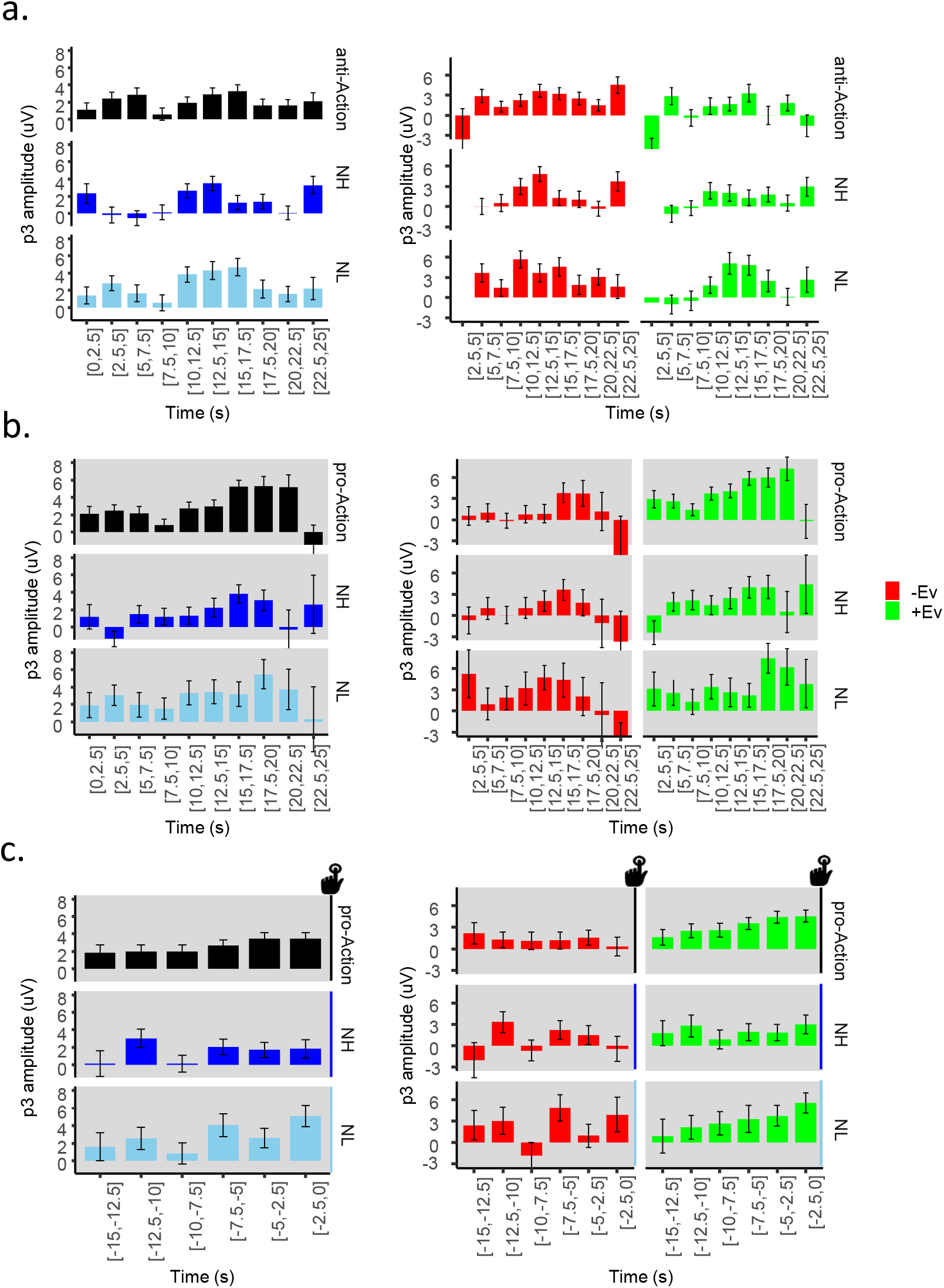
**A.** Stimulus-locked grand averaged (± SEM) P3 amplitude throughout trial time, in 2.5s bins collapsed (left) and for separate types of evidence (right) in No-Action trials. ***B.*** Stimulus-locked grand averaged (± SEM) P3 amplitude throughout trial time, in 2.5s bins collapsed (left) and for separate types of evidence (right) in Action trials ***C***. Action-locked grand averaged (± SEM) P3 amplitude throughout trial time, in 2.5s bins across (left) and for separate types of evidence (right) in Action trials.

**Figure S4.**
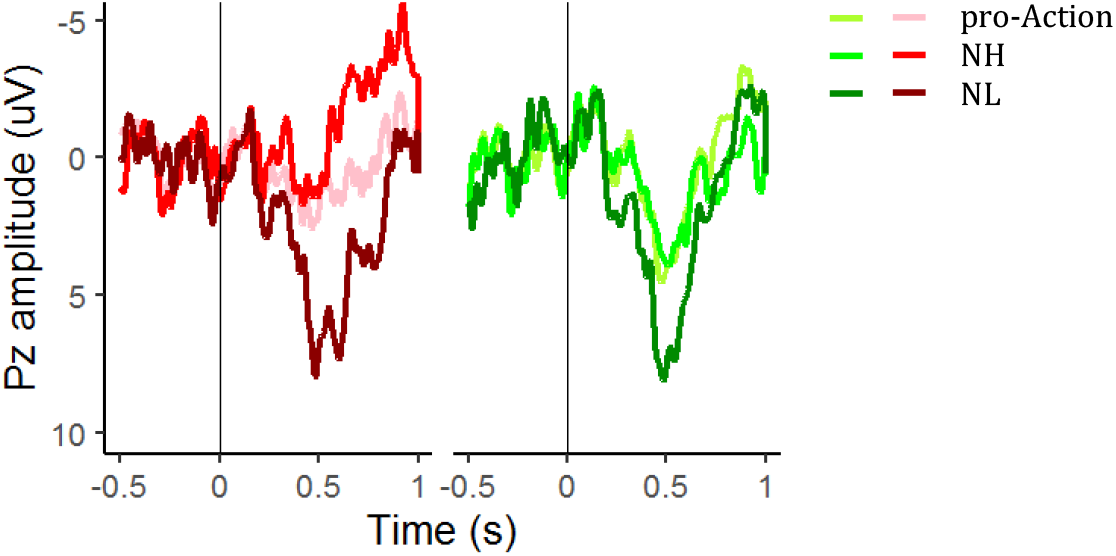
Grand-averaged ERP traces for the third +Ev (green) and –Ev (red) items presented after trial start. The P3 is clearly different in different conditions, suggesting that its amplitude does not *only* reflect the amount of accumulated evidence.

**Table S1.** Sequential P3 analysis

| **A. Full model:**<br>**P3 ~ EvidenceType\*Condition\*TimeToAction\*TimeToAction^2 + (1\|Sub/Block/Trial)** |  |  |  |
| --- | --- | --- | --- |
| **Fixed Factor** | **Chisq** | **Df** | **Pr(>Chisq)** |
| **TimeToAction** | 6.508264 | 1 | 0.010737 |
| **TimeToAction2** | 1.592671 | 1 | 0.206945 |
| **Condition** | 3.066541 | 2 | 0.215829 |
| **EvidenceType** | 14.46524 | 1 | 0.000143 |
| **TimeToAction:TimeToAction2** | 0.004932 | 1 | 0.944013 |
| **TimeToAction:Condition** | 0.624152 | 2 | 0.731926 |
| **TimeToAction2:Condition** | 0.95037 | 2 | 0.62177 |
| **TimeToAction:EvidenceType** | 3.985949 | 1 | 0.045881 |
| **TimeToAction2:EvidenceType** | 2.482823 | 1 | 0.115095 |
| **Condition:EvidenceType** | 2.086514 | 2 | 0.352305 |
| **TimeToAction:TimeToAction2:Condition** | 3.102155 | 2 | 0.212019 |
| **TimeToAction:TimeToAction2:EvidenceType** | 0.291098 | 1 | 0.589517 |
| **TimeToAction:Condition:EvidenceType** | 0.260738 | 2 | 0.877772 |
| **TimeToAction2:Condition:EvidenceType** | 0.546902 | 2 | 0.76075 |
| **TimeToAction:TimeToAction2:Condition:EvidenceType** | 3.74255 | 2 | 0.153927 |
| **B. Positive evidence (+Ev):**<br>**P3 ~ Condition\*TimeToAction\*TimeToAction^2 + (1\|Sub/Block/Trial)** |  |  |  |
| **Fixed Factor** | **Chisq** | **Df** | **Pr(>Chisq)** |
| **TimeToAction** | 10.20417 | 1 | 0.001401 |
| **TimeToAction2** | 4.093733 | 1 | 0.043042 |
| **Condition** | 3.605364 | 2 | 0.164856 |
| **TimeToAction:TimeToAction2** | 0.169652 | 1 | 0.680421 |
| **TimeToAction:Condition** | 0.858792 | 2 | 0.650902 |
| **TimeToAction2:Condition** | 0.862283 | 2 | 0.649767 |
| **TimeToAction:TimeToAction2:Condition** | 1.569641 | 2 | 0.456202 |
| **C. Negative evidence (-Ev):**<br>**P3 ~ Condition\*TimeToAction\*TimeToAction^2 + (1\|Sub/Block/Trial)** |  |  |  |
| **Fixed Factor** | **Chisq** | **Df** | **Pr(>Chisq)** |
| **TimeToAction** | 0.116999 | 1 | 0.732313 |
| **TimeToAction2** | 0.306222 | 1 | 0.580008 |
| **Condition** | 2.390738 | 2 | 0.302592 |
| **TimeToAction:TimeToAction2** | 0.093114 | 1 | 0.760255 |
| **TimeToAction:Condition** | 0.222649 | 2 | 0.894648 |
| **TimeToAction2:Condition** | 0.813593 | 2 | 0.66578 |
| **TimeToAction:TimeToAction2:Condition** | 4.557156 | 2 | 0.10243 |

**Table S2.** Trial-by-trial average p3 analysis

| **A. Full model:**<br>**P3 ~ EvidenceType\*Condition\*Decision + (1\|Sub/Block/Trial)** |  |  |  |
| --- | --- | --- | --- |
| **Fixed Factor** | **Chisq** | **Df** | **Pr(>Chisq)** |
| **EvidenceType** | 0.958937 | 1 | 0.327455 |
| **Condition** | 4.3484 | 2 | 0.113699 |
| **Decision** | 0.493595 | 1 | 0.482328 |
| **EvidenceType:Condition** | 0.539244 | 2 | 0.763668 |
| **EvidenceType:Decision** | 31.56472 | 1 | <0.001 |
| **Condition:Decision** | 0.817029 | 2 | 0.664637 |
| **EvidenceType:Condition:Decision** | 3.233602 | 2 | 0.198533 |
| **B. No-Action:**<br>**P3 ~ EvidenceType\*Condition + (1\|Sub/Block/Trial)** |  |  |  |
| **Fixed Factor** | **Chisq** | **Df** | **Pr(>Chisq)** |
| **EvidenceType** | 12.82875 | 1 | 0.000341 |
| **Condition** | 2.210082 | 2 | 0.331197 |
| **EvidenceType:Condition** | 0.418275 | 2 | 0.811284 |
| **C. Action:**<br>**P3 ~ EvidenceType\*Condition + (1\|Sub/Block/Trial)** |  |  |  |
| **Fixed Factor** | **Chisq** | **Df** | **Pr(>Chisq)** |
| **EvidenceType** | 18.12249 | 1 | <0.001 |
| **Condition** | 3.385639 | 2 | 0.184 |
| **EvidenceType:Condition** | 2.635856 | 2 | 0.267689 |

